# Documenting *Bombus nevadensis* in Minnesota, with some notes on discerning it from *B. auricomus* (Hymenoptera: Apidae)

**DOI:** 10.1101/2022.05.23.493105

**Authors:** Zachary M. Portman, Chan Dolan

## Abstract

In the face of well-documented declines in multiple bumblebee species, it is important to accurately identify species and properly delineate species ranges. Here, we document the range of *Bombus auricomus* (Robertson) and *B. nevadensis* Cresson in Minnesota, with particular reference to the unexpected discovery of *B. nevadensis* in St. Paul. We clarify the relative ranges of these two species and provide additional information on how to reliably identify them in Minnesota using color patterns and morphology, including differences in male genitalia. Our results support the consensus that *B. auricomus* and *B. nevadensis* are distinct species. Community science records were integral to fully documenting the range of *B. nevadensis* in Minnesota. Our findings demonstrate the value of community science data, though it highlights the need for experts to check the data and to be mindful of biases in observations around population centers.

## Introduction

The genus *Bombus*, despite being fairly well-resolved taxonomically, still has many issues. In the United States there are still new cryptic species being discovered, such as *B. bifarius* Cresson and *B. vancouverensis* Cresson (Ghisbain et al. 2020). Other species complexes remain unresolved, such as the *Bombus fervidus* complex (Koch et al. 2018). Even eastern species have taxonomic and identification issues and are difficult to identify, such as *B. vagans* Smith and *B. sandersoni* Franklin (Milam et al. 2020). Particularly in the face of recent declines in multiple bumblebee species (Cameron et al. 2011, Wood et al. 2019, Guzman et al. 2021), it’s important to properly delineate species boundaries and range extents.

The sister species *B. auricomus* (Robertson) and *B. nevadensis* Cresson are an example of species with a history of taxonomic uncertainty. The early taxonomic history of *B. auricomus* and *B. nevadensis* is relatively straightforward. *Bombus nevadensis* was described by Cresson (1874) from Nevada. *Bombus auricomus* was described (as *Bombias auricomus*) by Robertson (1903) from Illinois. It seems likely that Robertson was not familiar with *B. nevadensis* since he did not include it in the genus *Bombias*, which Robertson (1903) created to include *B. auricomus, B. fraternus* (Smith), and *B. griseocollis* (De Geer). However, the status of *B. auricomus* and *B. nevadensis* was clarified by Franklin (1913), who recognized both species as distinct and provided both morphological and color characters to separate them. Swenk (1907) did not have difficulty separating the two species in Nebraska, an area where both species overlap in range.

Later on, however, the range overlap and similar color patterns of *B. auricomus* and *B. nevadensis* caused confusion. Milliron (1961, 1971) and Webb and LaBerge (1962) considered *B. auricomus* and *B. nevadensis* a single species, and relegated *B. auricomus* to a subspecies of *B. nevadensis*. The name *Bombus nevadensis nevadensis* was used for the western form, and *B. n. auricomus* was used for the eastern form. The justification for this decision was based on overlapping color patterns as well as the claimed presence of hybrids (Laberge and Webb 1962, Milliron 1971). However, morphological characters, particularly the genitalia characters used by Franklin (1913), were not discussed in any of the justifications for grouping *B. auricomus* and *B. nevadensis* as a single species (Milliron 1961, 1971, Laberge and Webb 1962). This classification was subsequently accepted by most authors, including Mitchell (1962).

More recently, evidence has steadily accumulated to support *B. auricomus* and *B. nevadensis* as distinct species. Scholl et al. (1992) classified them as distinct species based on analysis using electrophoresis data. Their results suggested that overlapping color patterns between *B. auricomus* and *B. nevadensis* were merely convergence on the same Mullerian complexes rather than indicative of hybridization between the species (Scholl et al. 1992). This was further supported by morphological (Williams 1998) and genetic data (Cameron et al. 2007). As a result, the specific status of *B. auricomus* and *B. nevadensis* has become widely accepted (e.g. Colla et al. 2011, Williams et al. 2014). However, as a result of the taxonomic confusion, the exact extent of the range of the two species, particularly in areas where they overlap, is unclear. Further, there are difficulties in distinguishing the two species; the most recent identification resource is based on overlapping color patterns and a single subtle morphological character (Williams et al. 2014).

Here, we have three primary objectives. First, we report the unexpected discovery of *B. nevadensis* in St. Paul, Minnesota. Second, we use a combination of museum and community science records to understand the range of *B. nevadensis* in Minnesota and understand to what degree the ranges of *B. auricomus* and *B. nevadensis* overlap in the state. Third, we provide additional information on the identification of the two species. Our study highlights some of the promises and pitfalls of community science data, and we discuss the utility of this data for documenting bumblebees and contributing to scientific investigations.

## Methods & Materials

For this study we examined a combination of museum specimens, specimens from recent studies, and iNaturalist observations. We reexamined all available pinned specimens of *Bombus auricomus* in Minnesota. The source of these specimens were the University of Minnesota Insect Collection (UMSP, ∼219 specimens), the Minnesota Department of Natural Resources (76 specimens), recent projects from the Cariveau lab (18 specimens, Lane et al. (2020, 2022); 1 specimen, B. Bruninga-Socolar, unpublished; 12 specimens, D. Cariveau, unpublished), the Spivak lab (1 specimen, Wolfin et al. (2021)), unpublished data from Elaine Evans (34 specimens), a recent thesis on the bees of western Minnesota (12 specimens, Pennarola (2019)), and a study on the bees of Six Mile Marsh (1 specimen, 30 observations, Portman et al. In Press). In total, approximately 362 Minnesota specimens of *Bombus auricomus* were examined. We also examined 12 specimens (11 females and 1 male) of *Bombus nevadensis* from the University of Minnesota Insect Collection from throughout the range of that species (Colorado, Nevada, New Mexico, South Dakota, Utah, and Washington).

We supplemented the traditional museum and project specimens with an in-depth examination of all of the Minnesota observations of *B. auricomus* and *B. nevadensis* on iNaturalist (accessed 8 Sep 2021). In total, 831 observations of Minnesota *B. auricomus* or *B. nevadensis* were made or confirmed by ZP. Only observations that were personally confirmed by ZP were included in the study and maps. Finally, photo observations of *B. nevadensis* were supplemented from other sources, including Twitter (1 observation), and surveys by the Minnesota Department of Natural Resources (2 observations).

Spatial datasets were derived from museum specimens and confirmed observations on iNaturalist. Where necessary, historic museum specimens of *B. auricomus* and *B. nevadensis* in Minnesota were georeferenced using Google Earth Pro (v7.3.4.8248) to obtain latitude and longitude coordinate points. Observations of *B. auricomus* and *B. nevadensis* were downloaded directly from iNaturalist. Any iNaturalist observations with obscured coordinates were excluded from mapping. Figures were made using the open-source program QGIS (v3.16.2).

Morphological terminology follows Michener (2007), including the terminology for male terminalia (see Fig. 119-3 from Michener (2007)). Specimen images were taken using an Olympus DP27 camera mounted on an Olympus SZX16 stereo microscope, and photos were stacked using CombineZP software (Hadley 2010). Figures were made with Adobe Photoshop 2018 software (Adobe Systems Inc., San Jose, CA).

## Results

### St. Paul observations

In 2020 and 2021, ZP made four separate sightings of *Bombus nevadensis* in St. Paul, Ramsey Co., Minnesota. The details of each observation are:

1. 23 Jul 2020, a worker was spotted by ZP foraging on *Monarda fistulosa* in Horton Park (44.9640, -93.1572). Photographs were taken of the bee but it was not collected (Fig. 1A). The wings of the bee were totally unworn.
2. 25 Jul 2020, a worker foraging on *Monarda fistulosa* at the same location as the first sighting. Photographs were taken of the bee but it was not collected (Fig. 1B). The wings of the bee were heavily worn. This bee was likely not the same as the first observation since it was seen only two days later and much more worn.
3. 4 August 2020, a worker foraging on *Monarda didyma* in a residential garden less than a block from Horton Park. The specimen was collected after taking a few photographs (Fig. 1C). The wings of the bee were moderately worn. This bee is not the same as the second observation, since the wings were less worn, though it could potentially be the same bee as the first observation.
4. 4 May 2021, a queen foraging on a crabapple tree (*Malus* sp.) on a front lawn in a residential area one block from Horton Park (44.9630, -93.1605). Photographs were taken but it was not collected (Fig. 1D).

**Figure 1.**
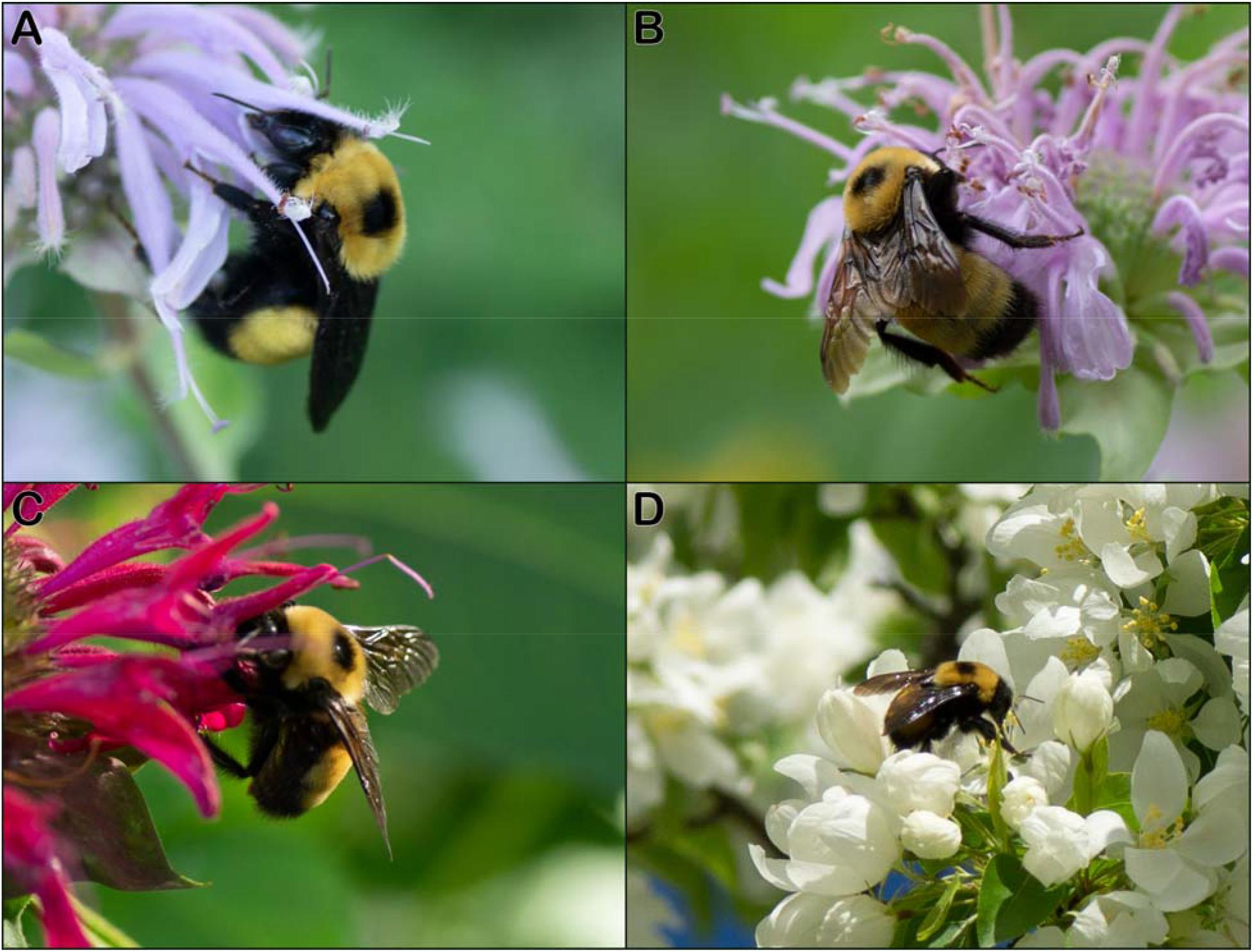
Observations of *Bombus nevadensis* in St. Paul, MN: (A) Worker sighted on 23 Jul 2020 in Horton park; (B) Worker sighted on 25 Jul 2020 in Horton Park; (C) Worker sighted and collected on 4 Aug 2020 near Horton Park; (D) Queen sighted on 4 May 2021 near Horton Park.

Overall, there were four sightings representing at least three individuals. One specimen (from the third observation) was collected and confirmed to match *B. nevadensis*. It is deposited in the Cariveau Native Bee Lab synoptic collection.

### *Reexamination of Minnesota specimens of* Bombus auricomus

The observations of *B. nevadensis* in St. Paul prompted a thorough review of the *B. auricomus* specimens in the UMSP and Cariveau Lab collections. Over 360 specimens were examined, and two were determined to actually be *B. nevadensis* rather than *B. auricomus*.

Both specimens were from a study on prairie restorations in western Minnesota by Pennarola (2019), which ZP had originally misidentified as *B. auricomus* when he identified them in 2018. The two specimens are:

1. 18 Jun 2016, female (worker), Yellow Medicine Co. (44.7072, -96.3972), Pennarola leg., net, *Trifolium pratense* [USGSDRO490051].
2. 10 Jul 2016, female (worker), Ulen WMA, Clay Co. (47.0820, -96.3382), Pennarola & Leone leg., bowl trap [USGSDRO508404].

### Community science records from iNaturalist

There have been 10 total observations of *B. nevadensis* in Minnesota posted to the community science website iNaturalist. Some records may be observations of the same individual.

1. 8 Aug 2016, queen, Lake Harbor, Lake Co., observed by Melissa Rainville on iNaturalist (username elissrainville, https://www.inaturalist.org/observations/3842280).
2. 31 Aug 2016, queen, Duluth, St. Louis Co, observed by iNaturalist user icenine5580 (https://www.inaturalist.org/observations/4006167)
3. 5 Aug 2018, male, Hawk Ridge Bird Observatory, St. Louis Co., observed by iNaturalist user dexternienhaus (https://www.inaturalist.org/observations/23094192).
4. 8 Aug 2019, male, Duluth, St. Louis Co., observed by iNaturalist user snapdragon822 (https://www.inaturalist.org/observations/30450704).
5. 25 Jul 2020, female (queen?), Duluth, St. Louis Co., observed by iNaturalist user dssmn (https://www.inaturalist.org/observations/54306526).
6. 26 Jul 2020, female (queen?), Duluth, St. Louis Co., observed by iNaturalist user dssmn (https://www.inaturalist.org/observations/54461388).
7. 26 Jul 2020, female (queen?), Duluth, St. Louis Co., observed by iNaturalist user dssmn (https://www.inaturalist.org/observations/54461740). Observed six minutes after the previous observation and may be the same bee.
8. 5 Aug 2020, male, Duluth, St. Louis Co., observed by iNaturalist user davidenrique (https://www.inaturalist.org/observations/55619527).
9. 27 Jul 2020, worker, Duluth, St. Louis Co, observed by iNaturalist user dssmn (https://www.inaturalist.org/observations/54519238).
10. 30 Jul 2021, worker, Duluth, St. Louis Co., observed by Tina Boucher on iNaturalist (username tina_boucher, https://www.inaturalist.org/observations/89237669).

### Other community scientist records

One observation of *B. nevadensis* was posted to Twitter.

1. June 2021, female (queen?), Warroad, Roseau Co., observed by Twitter user @Hogan698 (https://twitter.com/Hogan698/status/1402330272641728515).

### Observations from the Minnesota Department of Natural Resources

Two observations of *B. nevadensis* were provided from bee surveys by the Minnesota Department of Natural Resources. Specimens were not collected but photos were taken and confirmed.

1. 2 Aug 2021, female, Murray Co. (43.915525, -95.968497), observed by Lisa Gelvin-Innvaer of the Minnesota Department of Natural Resources, foraging on *Monarda fistulosa*. Photos confirmed by ZP.
2. 3 Jun 2021, queen, Lincoln Co.(44.2643, -96.3083), observed by Bob Dunlap of the Minnesota Department of Natural Resources, foraging on *Hydrophyllum virginianum*. An online record of the observation is available at https://www.bumblebeewatch.org/app/#/bees/view/85087.

### Summary and distribution of Bombus auricomus and B. nevadensis

In total, there are 19 records of *B. nevadensis* in MN: 4 sightings by ZP, 2 records from MNDNR surveys, 2 museum specimens from the University of Minnesota Insect Collection, 10 iNaturalist observations, and 1 community science observation from Twitter. In contrast, *B. auricomus* is much more common and abundant in Minnesota, with over 1240 records of *B. auricomus* (822 iNaturalist observations, 391 collected specimens, and 30 other observations). There were more iNaturalist observations than records from traditional methods, though some of the iNaturalist observations may be the same bee (e.g. iNaturalist observation 7 of *B. nevadensis* was observed six minutes before observation 6 by the same user at the same location). In addition, iNaturalist observations are generally concentrated around population centers (Fig. 2B).

**Figure 2.**
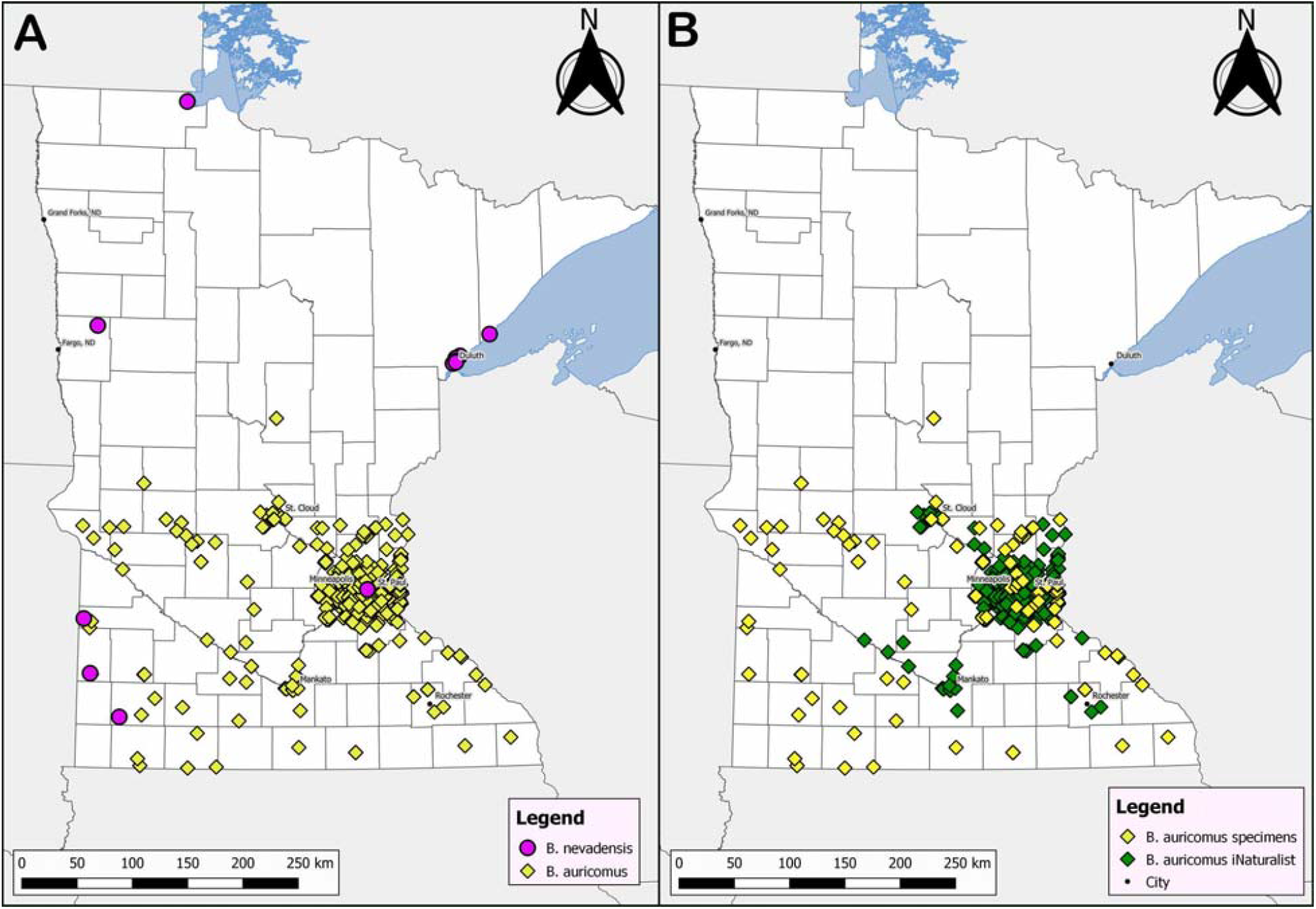
(A) Ranges of *Bombus nevadensis* (purple dots) and *B. auricomus* (yellow dots) in Minnesota; (B) Breakdown of *B. auricomus* records between museum specimens (yellow dots) and community science observations (green dots).

We found that there is overlap between *B. auricomus* and *B. nevadensis* in some areas of the state but not others. *Bombus auricomus* primarily occurs in the southern half of the state, whereas *B. nevadensis* primarily occurs in the northern and western areas of the state (Fig. 2A).

We were unable to confirm any northern records of *B. auricomus* in Minnesota, and as a result, all northern records of the subgenus *Bombias* appear to be *B. nevadensis. Bombus auricomus* and *B. nevadensis* overlap in two areas of the state: southwestern areas of Minnesota and in St. Paul (Fig. 2A).

### *Identification of* Bombus auricomus *and* Bombus nevadensis *in Minnesota*

Examination of available material of *Bombus auricomus* and *B. nevadensis* revealed both color and morphological characters that allow for their consistent identification in Minnesota and surrounding states. Based on the relatively sparse male material, identification of males based on color is less certain, but they can be separated by the genitalia.

Female *B. auricomus* and *B. nevadensis* share the same basic color pattern of having the dorsum of the thorax yellow in part, the sides of the thorax black, T1 partially yellow, and T2−3 yellow, with the rest of the abdomen black. However, *B. auricomus* always have a complete black band between the wing bases, whereas *B. nevadensis* have the black hairs limited to a black square medially (Fig. 3B), and the scutum hairs can even be entirely yellow. Occasionally *B. nevadensis* have dark hairs that extend all the way to the tegula, but these are always intermixed with the light hairs. In contrast, *B. auricomus* always has the dark hairs extend to the tegula (Fig. 3A) with at most a few intermixed light hairs.

**Figure 3.**
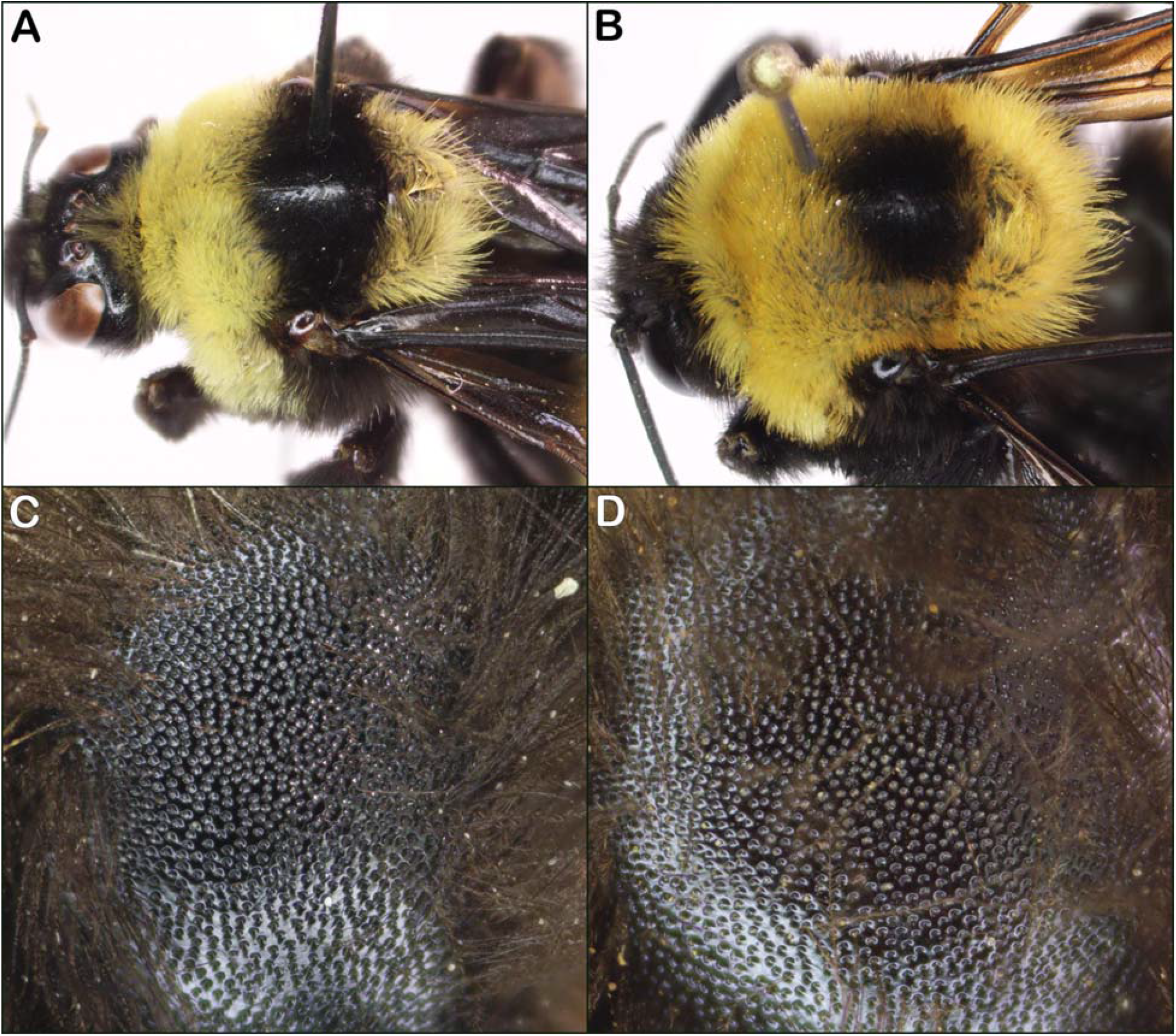
(A) Light worker of *Bombus auricomus*; (B) Typical color pattern of *Bombus nevadensis*; (C) Punctures on the side of *B. auricomus*, showing dense punctures; (D) Punctures on the side of *B. nevadensis*, with punctures separated by about 1 puncture width.

*Bombus auricomus* often has black hairs on the scutellum, whereas *B. nevadensis* has the scutellum entirely yellow (Fig. 3B). In lighter specimens of *B. auricomus* (as in Fig. 3A), the scutellum is fully yellow, but in these cases, the hairs on the vertex are also predominantly yellow (Fig. 3A), whereas *B. nevadensis* always has mostly black hairs on the vertex (as in Fig. 4C). On the abdomen, *B. auricomus* typically have the hairs on T1 predominantly black, whereas *B. nevadensis* have the hairs on T1 predominantly yellow, though this is variable.

**Figure 4.**
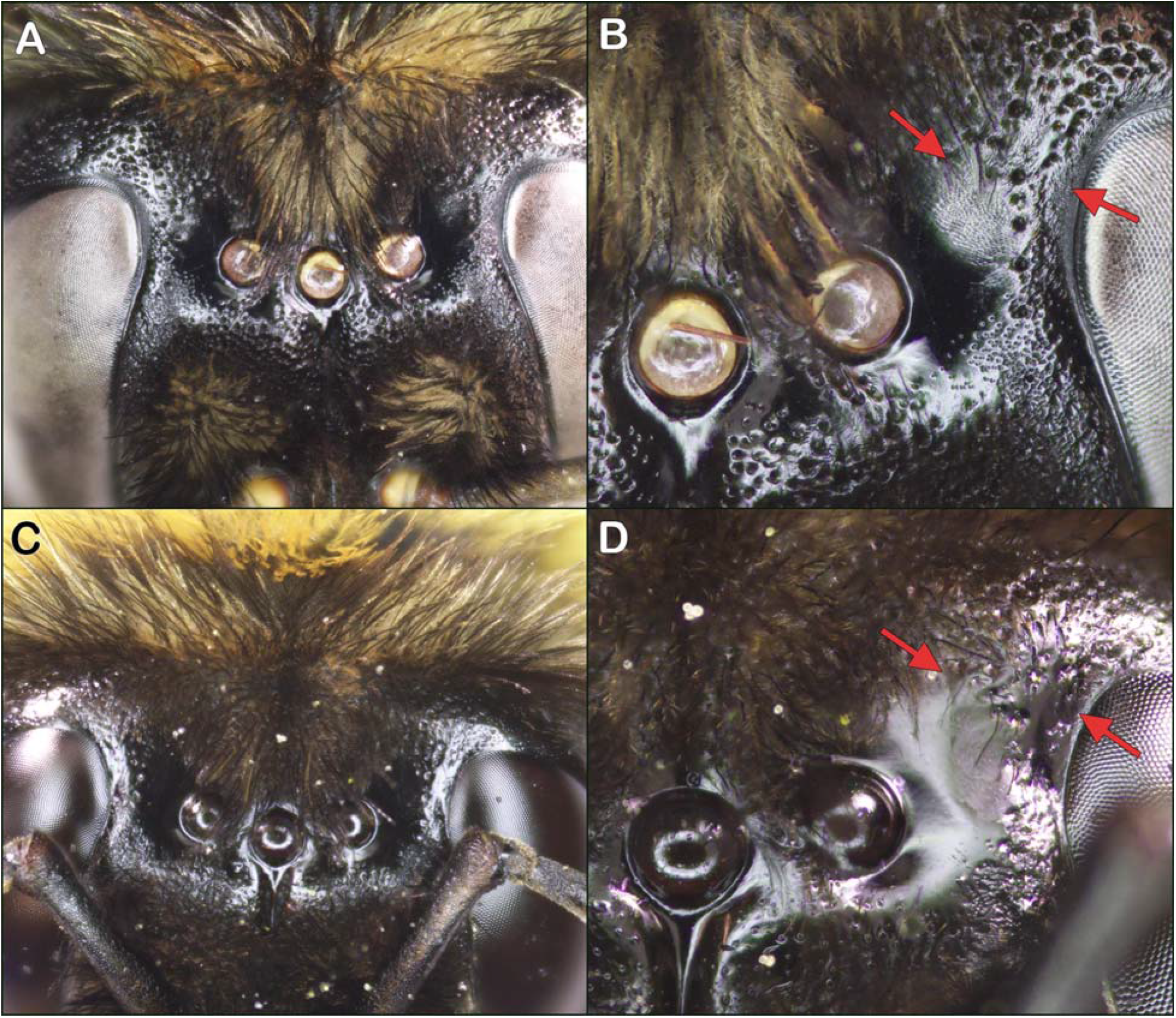
Sculpturing around the occelli of *Bombus auricomus* and *B. nevadensis*: (A) *Bombus auricomus* ocellar area; (B) Zoomed-in view of *B. auricomus* ocellar area, arrows indicate areas with tessellation; (C) *B. nevadensis* ocellar area; (D) Zoomed in view of *B. nevadensis* ocellar area, arrows indicate smooth areas that lack tessellation.

Finally, there are subtle but consistent differences in the shade of the yellow hairs. *Bombus auricomus* are more of a pale, whitish-yellow, whereas *B. nevadensis* are more of a darker yellow. However, fading of hairs makes this character unreliable.

In addition to color characters, females can also be separated using morphological characters. Previous researchers have pointed to the sculpturing of the area lateral to the ocelli as a diagnostic character for splitting auricomus and nevadensis (Williams 1998, Williams et al. 2014). Specifically, the area between the ocelli and the eye is tessellate in *B. auricomus* (Figs 4A, B) but is mirror-smooth in *B. nevadensis* (Figs 4C, D). Our own examination has largely supported this character, though we have found two *B. auricomus* that lack tessellation in this area. In addition, the punctures of *B. auricomus* on the face and thorax are consistently larger and coarser (Figs 3C, 4B), whereas *B. nevadensis* have the punctures finer and more separated (Figs 3D, 4D). This character is subtle but consistent, and is most apparently on the sides, scutum, and vertex.

Male *B. auricomus* and *B. nevadensis* are more difficult to separate than females. Both species are characterized by their greatly enlarged eyes and both typically have T1−3 yellow and a primarily yellow thorax. In general, *B. nevadensis* males have more extensive lighter coloration than *B. auricomus*. Specifically, *B. nevadensis* typically have the thorax entirely yellow, T4 yellow, at least in part, and orange hairs on T5 and T6. In comparison, *B. auricomus* typically have T4 black, have at least some black on the scutum, and T5 and T6 have only black hairs. The presence of orange hairs at the apex of the abdomen in *B. nevadensis* has typically been treated as diagnostic. However, in the UMSP collection, there are some *B. auricomus* males with a small amount of orange hairs on the tip of the abdomen as well. As a result, a bee with T1−3 yellow and with a small amount of orange on T5−6 can potentially be either *B. nevadensis* or *B. auricomus*. Finally, the extent of black hairs on the dorsum of the thorax is extremely variable in *B. auricomus*, ranging from an entire black band to entirely yellow.

Morphologically, we have only found genitalia characters that can split *B. auricomus* and *B. nevadensis* males. Previous researchers have pointed to the tessellation around the ocelli as a potential splitting character for males (Williams et al. 2014), but we have not been able to discern that character in our lone available male specimen. Additional material may reveal external morphological differences. Although we were not able to find consistent external morphological differences, the genitalia can separate *B. auricomus* and *B. nevadensis*. Specifically, the apex of the gonostylus is more narrowed and pointed inwards in *B. auricomus* (Fig. 5A, little red arrow), which results in a broader lateral flap of the volsella along the border of the gonostylus (Figs 5A, C, big red arrow). In comparison, *B. nevadensis* has the gonostylus more broadly rounded (Fig. 5B, little red arrow), with a narrower extension of the volsella along the lateral margin of the gonostylus (Figs 5B, D, big red arrow). These genitalia characters were previously identified and illustrated by Franklin (1913, reproduced in Fig. 6), though we have not found mention of this character by any subsequent authors. Additional genitalia characters include the shape of the apicomedial margin of the gonocoxite, which is more rounded in *B. auricomus* (Fig. 5A, 6A) and more squared-off in *B. nevadensis* (Fig. 5B, 6B). In addition, *B. auricomus* has the penis valves noticeably thicker when viewed laterally (Fig. 5C) compared to *B. nevadensis* (Fig 5D).

**Figure 5.**
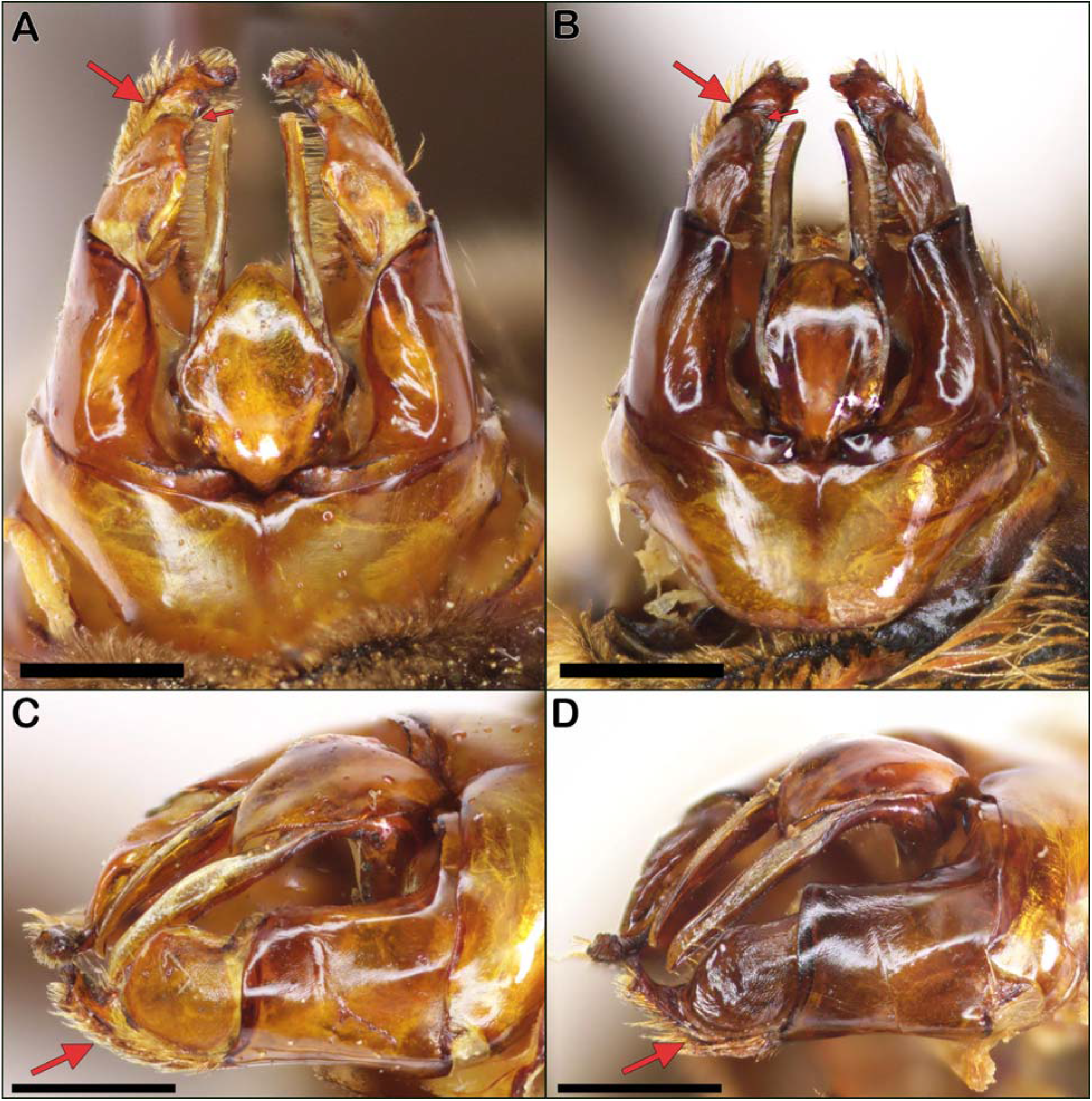
(A) *Bombus auricomus* dorsal view; (B) *B. nevadensis* dorsal view; (C) *B. auricomus* lateral view; (D) *B. nevadensis* lateral view. All scale bars = 1 mm.

**Figure 6.**
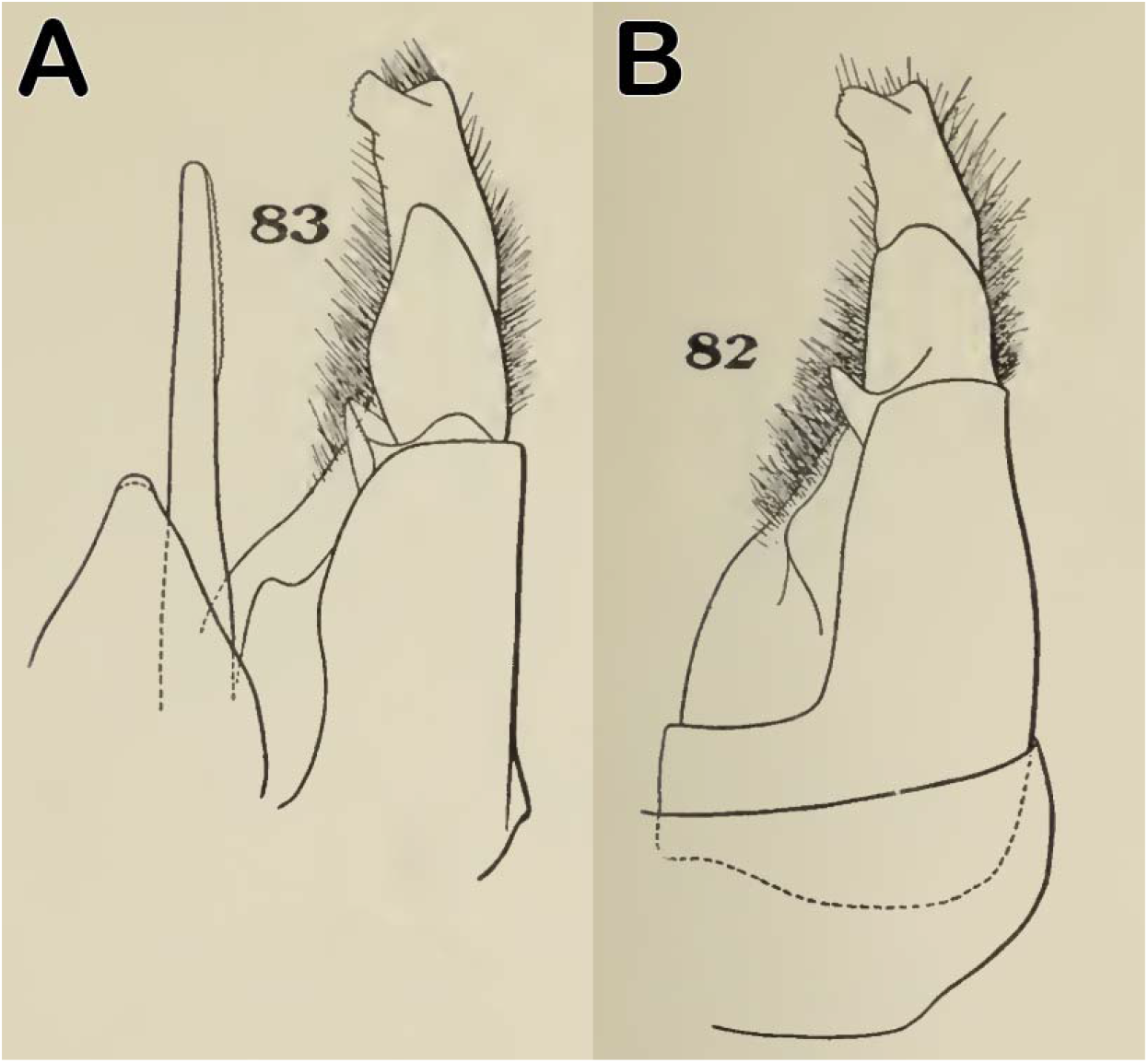
Illustrations of the dorsal genitalia adapted from Franklin (1913): (A) *Bombus auricomus*; (B) *Bombus nevadensis*.

## Discussion

Our findings demonstrate that both *Bombus auricomus* and *B. nevadensis* occur in Minnesota. This represents an expansion of the known range of *B. nevadensis*, which previously had its easternmost record in eastern North Dakota (Williams et al. 2014). Despite the lack of records prior to 2016, it seems clear that Minnesota is part of the historic range of *B. nevadensis*. Particularly in the western areas of the state, these areas are close enough to the known historic range of *B. nevadensis* in North Dakota and Manitoba (Williams et al. 2014) that their absence can likely be chalked up to a lack of historic sampling in those areas. In addition, the number of observations of *B. nevadensis* in the Duluth area suggests that the bee has been there for some time rather than being a recently established population. That same area along Lake Superior also hosts populations of another primarily western bumblebee, *B. melanopygus* Nylander, and similar historic biogeographic factors may be influencing both species. Further, we only examined Minnesota specimens, and given the tangled taxonomic history between *B. auricomus* and *B. nevadensis*, it is possible that historic specimens of *B. nevadensis* exist in collections but have been misidentified. Finally, given the range of *B. nevadensis* in Minnesota, our results raise the possibility that *B. nevadensis* occurs in western Wisconsin and northwestern Iowa.

Our discovery of *B. nevadensis* in St. Paul was unexpected. Although it is likely that western and northern Minnesota is part of the historic range of *B. nevadensis*, the population in St. Paul has a more uncertain provenance. Based on the observation of *B. nevadensis* in St. Paul over two years (three workers in 2020 and one queen in 2021) it indicates that there is (or was) an established and breeding population of *B. nevadensis* present. Given the lack of historic records of *B. nevadensis* in the area, combined with the relatively intensive sampling and observations in and around the Twin Cities, this suggests that the St. Paul *B. nevadensis* population has established relatively recently. However, this is relatively speculative. More work is needed to determine the extent of the St. Paul population and to see if it persists. If the St. Paul population is new, it would join a few other US bumblebee species that are expanding their range, such as *B. impatiens* Cresson and *B. bimaculatus* Cresson (Colla et al. 2012, Jacobson et al. 2018, Guzman et al. 2021).

Our findings agree with the consensus that *B. auricomus* and *B. nevadensis* are distinct species (Scholl et al. 1992, Williams 1998, Cameron et al. 2007, Colla et al. 2011, Williams et al. 2014). In particular, differences in the morphology of the genitalia—originally documented by Franklin (1913) and expanded upon here—provide definitive support for this hypothesis. That these differences in genitalia were not discussed by subsequent workers who synonymized the species (e.g. Webb and LaBerge 1962, Milliron 1971) is surprising, and it demonstrates the value of reviewing older taxonomic literature, which can contain insights and high-quality work. Due to lack of material we have been unable to find external morphological differences in males, though these may be found in the future with the examination of more specimens. In addition, more work is needed to explore and document the geographic patterns of color patterns in both males and females, since the color patterns of *B. auricomus* and *B. nevadensis* are both variable and overlapping.

Our understanding of *B. nevadensis* in Minnesota benefitted from the community science platform iNaturalist. Community science is a powerful tool for generating large amounts of data but it has certain drawbacks (Silvertown 2009, Theobald et al. 2015). Particularly in bees, the observations generally require expert confirmation (Falk 2019, MacPhail et al. 2020), which was the case in our study, with many of the first observations of *B. nevadensis* misidentified as *B. auricomus*. Further, data collected by community scientists are mostly concentrated in urban areas and high population centers (Geldmann et al. 2016). This can lead to a sampling bias, where rural areas are relatively under surveyed using community science methods. This bias is evident when comparing the distribution of *B. auricomus* from community science observations on iNaturalist and entomological specimens collected by researchers (Fig. 2B). Observations collected from iNaturalist are mostly concentrated in eastern Minnesota, with most observations being in the Minneapolis-Saint Paul metropolitan area. In contrast, specimens collected by researchers were often collected from areas where there are no observations of *B. auricomus* via iNaturalist. A similar pattern was seen with the four westernmost records of *B. nevadensis*, which were found during surveys by researchers at the University of Minnesota and the Minnesota Department of Natural Resources. This comparison shows the strength of comunity science as a means for data collection in population centers but also the importance of accounting for bias when using community-science data (Geldmann et al. 2016). If the use of volunteer-based data collection websites is continued to be used in monitoring, expanding usage outside of urban areas will be an important next step.

## Acknowledgments

We thank Lisa Gelvin-Innvaer and Bob Dunlap of the Minnesota Department of Natural Resources for sharing information and photographs of their observations of *Bombus nevadensis*, and we thank Joel Neylon, Christopher Smith, and Tony Ernst for helping find *B. nevadensis* observations. We also thank UMSP curator Robin Thomson for loaning *Bombus* specimens.

